# Membrane surface geometry is a determinant of mitochondrial electron transfer and cellular adaptation

**DOI:** 10.64898/2026.09.12.751109

**Authors:** Miguel González-Hernández, Carmen Choya-Foces, Daymi Céspedes de los Ríos, Enrique Calvo, José Luis Cabrera-Alarcón, Cristián Huck-Iriart, Rebeca Acín-Pérez, Judith Langer, John W Elrod, Jesús Vázquez, Jesús Ruiz-Cabello, Susana Carregal-Romero, Pablo Hernansanz-Agustín

**Affiliations:** Centro de Neurociencias Cajal (CNC-CSIC), Madrid, Spain; Aging + Cardiovascular Discovery Center, Lewis Katz School of Medicine, Temple University, Philadelphia, PA, USA; Unidad de Investigación, Hospital Universitario Santa Cristina, Instituto de Investigación Sanitaria Princesa (IIS-IP), Madrid, Spain; Centro Nacional de Investigaciones Cardiovasculares Carlos III (CNIC), Madrid, Spain; CIBER de Enfermedades Cardiovasculares (CIBERCV), Madrid 28029, Spain; Centro de Investigación Biomédica en Red de Fragilidad y Envejecimiento Saludable, Spain; ALBA Synchrotron Light Source, 08290 Cerdanyola del Vallès, Spain; Center for Cooperative Research in Biomaterials (CIC biomaGUNE), Basque Research and Technology Alliance (BRTA), Donostia San Sebastián, Spain; Centro de Investigación Biomédica en Red de Enfermedades Respiratorias, Spain; NMR and Imaging in Biomedicine Group, Department of Chemistry in Pharmaceutical Sciences, Pharmacy School, University Complutense Madrid, 28040 Madrid, Spain; Ikerbasque, Basque Foundation for Science, Bilbao, Spain

## Abstract

Energy conversion in living organisms relies on biological membranes that facilitate electron transfer between oxidoreductases. In mitochondria, this process is mediated by the electron transport chain embedded in the inner mitochondrial membrane (IMM). Under various physiological and genetic conditions, mitochondrial matrix Na⁺ levels increase, reducing IMM fluidity through the formation of ternary coordination adducts between Na⁺ and phospholipids. These adducts impair ubiquinone (CoQ) transfer between respiratory complexes, thereby promoting mitochondrial reactive oxygen species (mtROS) production and activating the hypoxic adaptive pathway. Here, we show that modifying solely the ionic subatomic interaction with phospholipids is sufficient to prevent initiation of this pathway. Compound A (CA) outcompetes Na⁺ for phospholipid binding without impairing CoQ transfer, thereby preventing mtROS production and hypoxic adaptation. This divergence arises from the penta-coordinate complexes formed by CA with phospholipids, in contrast to the trigonal adducts formed by Na⁺. This structural distinction preserves IMM fluidity because CA–phospholipid assemblies adopt a less angular configuration. These findings establish membrane-surface geometry, modulated by ion–phospholipid interactions, as an unexpected determinant of membrane biology, mitochondrial energy conversion, redox signalling, and cellular adaptation, with profound implications for physiology and disease.

## Main text

Energy conversion relies on the ability of biological membranes to enable the transfer of electron carriers between oxidoreductase enzymes. In eukaryotes, the IMM contains the electron transport chain, which coordinates the function of multiple complexes and supercomplexes to the electron transfer among them. Mitochondrial complexes I (CI) and II (CII) oxidize NADH and succinate, respectively, to reduce CoQ. Subsequently, complex III (CIII) utilizes this CoQ to reduce cytochrome c (cyt c), while complex IV (CIV) oxidizes cyt c, facilitating the reduction of O₂ to H₂O. These sequential redox reactions are coupled to the ejection of H⁺ to the intermembrane space by CI, CIII, and CIV, and to the Na^+^/H^+^ exchange (NHE) by CI^1^, across the IMM generating a H^+^ motive force (Δp). This electrochemical potential gradient drives the electrophoretic re-entry of H^+^ into the mitochondrial matrix, which is coupled to the phosphorylation of adenosine diphosphate (ADP) to adenosine triphosphate (ATP) via a fifth complex (CV), thus completing energy conversion in a process named oxidative phosphorylation (OxPhos).

Point mutations in the mitochondrial DNA (mtDNA) produce defects in OxPhos function. Human G11778A mutation translates into an Arg-to-His substitution in the ND4 gene of CI, which induces a defect in the NHE by this complex^1^. To understand the pathophysiology of this mutation, we sought to identify the molecular pathways activated in this genetic model. For that, we performed Gene Set Enrichment Analysis (GSEA) of proteomics data obtained from G11778A cybrid samples and compared it to their isogenic controls (Ctrl). G11778A showed a conspicuous enrichment of pathways related to the adaptation to decreased oxygen levels (Fig. 1a). In particular, G11778A was particularly abundant in hypoxia inducible factor (HIF)-1 target gene expression (Extended Data Fig. 1a and b), which correlated with a marked acidification by G11778A cells (Extended Data Fig. 1c).

**Figure 1.**
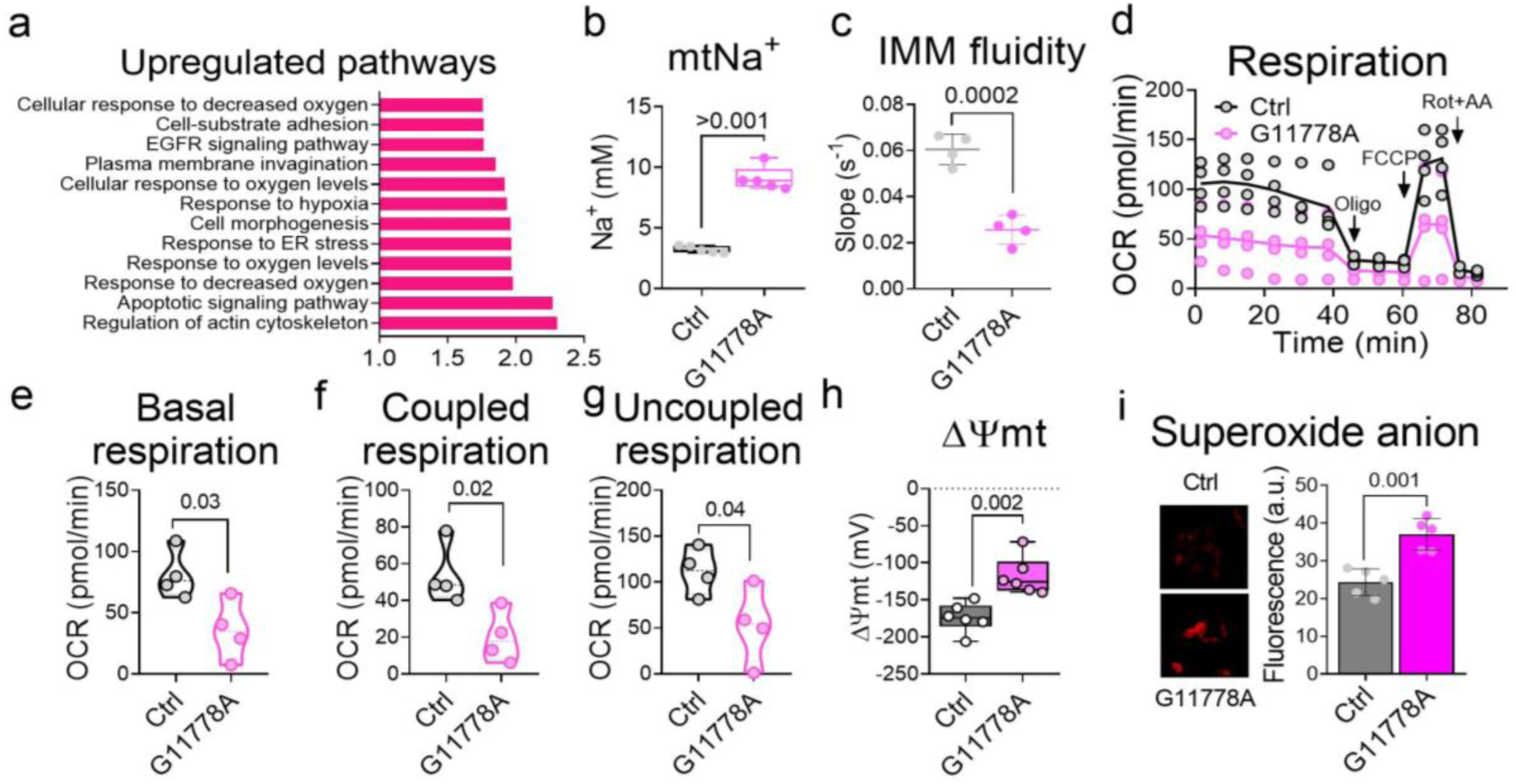
G11778A mutant phenotype is compatible with a constitute activation of the Na^+^-dependent ROS production mechanism. **(a)** Gene Set Enrichment Analysis (GSEA) showing the upregulated pathways in G11778A in comparison to its isogenic control (Ctrl; n=3). **(b)** Quantification of matrix Na^+^ by confocal microscopy in G11778A and Ctrl. **(c)** Quantification of FRAP of G11778A and Ctrl. **(d)** Oxygen consumption of G11778A Ctrl measured by Seahorse before and after the addition of 5 μM oligomycin (Oligo), 300 nM FCCP and 1 μM rotenone (Rot) and 1 μM Antimycin A (AA). **(e)** Basal respiration of G11778A and its isogenic control (Ctrl) measured by Seahorse. **(f)** Coupled respiration of G11778A and its isogenic control (Ctrl) measured by Seahorse. **(g)** Reserve capacity of G11778A and its isogenic control (Ctrl) measured by Seahorse. **(h)** Measurement of ΔΨmt in G11778A and Ctrl by confocal microscopy. **(f)** Measurement of O_2_^•-^ levels in G11778A and Ctrl by confocal microscopy. Left: Representative images showing fluorescence intensity; right: Quantification of fluorescence. Data are presented as mean ± percentiles in (b, c and e), mean in (d) and mean ± s.d. in (f). Unpaired two-tailed Student’s t-test was used in (b, c, e and f).

HIF-1 is transcriptional factor composed by a α and a β subunit^2^. Their activity is mainly regulated by the stability of their α subunits which is, in turn, regulated by the levels of O_2_^3^. Reactive oxygen species (ROS) are among the molecules able to promote the stabilization of HIFs α subunits^4–7^. ROS levels can be elevated after the accumulation of mitochondrial Na^+^ in a panoply of (patho)physiological conditions^8–12^. This cation interacts with phospholipids in the IMM, decreasing its fluidity, lowering electron transfer, specifically between CII and CIII, and promoting the production of superoxide anion (O_2_^•-^) by the latter^12^. Given that the G11778A mutant shows a defective mitochondrial Na^+^ exit capacity and that the accumulation of Na^+^ in the matrix promotes the production of ROS by CIII, we hypothesized that G11778A mutant may exhibit constitutive activation of the Na^+^-hindered electron transfer mechanism, which could be responsible for the activation of HIF-1. G11778A showed increased levels of mitochondrial Na^+^ and lowered IMM fluidity (Fig. 1b and c), which was associated to a decrease in respiration (Fig. 1d-f and Extended Data Fig. 1e-g). This occurred even in the absence of Δp, which is the major regulator of respiration (Fig. 1g), indicating that electron transfer in G11778A is hindered likely as a consequence of reduced IMM fluidity. On the one hand, this turned into mitochondrial depolarization (Fig. 1h). On the other, hampered IMM electron transfer correlated with an increased production of ROS (Fig. 1i), which were detectable despite the higher presence of a collection of antioxidant enzymes in the mutant (Extended Data Fig. 2a-e). This led to a downregulation of pathways involved in cell division (Extended Data Fig. 2f) and decreased cell proliferation (Extended Data Fig. 2g). These results indicate that G11778A exhibits a phenotype compatible with a constitutive activation of the Na^+^-dependent ROS production mechanism, which is based on the interaction of mitochondrial Na^+^ with phospholipids in the IMM, reducing its fluidity, hampering mitochondrial electron transfer, increasing ROS generation and activating the hypoxic adaptive pathway.

As Na^+^ is able to interact with phospholipids in the mitochondria and reduce IMM fluidity, we hypothesised that a positively charged cation with a larger charge density would exert a stronger effect. For that, we compared the effects of Na^+^ with those of CA on mitochondrial membrane fluidity. In a first series of experiments, we measured the fluidity of isolated mitochondrial membranes in the absence or presence of NaCl or CA, from 4°C to 37°C. In this approach, generalized polarization (GP) values are inversely proportional to membrane fluidity. Unexpectedly, CA, in contrast to Na^+^, did not have any measurable effect either on the fluidity of mitochondrial membranes (Fig. 2a-b) or in their phase transition (Extended Data Fig. 3a), showing similar values than untreated membranes. We aimed to confirm these results using an alternative method. In this assay, a fluorescent probe was incorporated into mitochondrial membranes, which were subsequently divided into three samples. Probe release (i.e., decrease in fluorescence) was then monitored in the absence or presence NaCl or CA. We observed that, whereas NaCl interfered with probe release, CA-treated and untreated mitochondrial membranes behaved similarly, facilitating its scape (Fig. 2c and Extended Data Fig. 3c). This corroborated that CA, in contrast to Na^+^, was unable to reduce mitochondrial membrane fluidity. In a second series of experiments, we assayed the composed CII+CIII activity in isolated mitochondrial membranes in the absence or presence of NaCl and CA. In conditions in which individual CII and CIII activities remain unchanged, electron transfer between these complexes is a functional readout of IMM fluidity. Strikingly, while NaCl reduced CII+CIII, CA was not able to modify it (Fig. 2d-f). These results indicate that, despite the larger charge density of CA, this cation does not have the same effect as Na^+^ on IMM fluidity and electron transfer.

**Figure 2.**
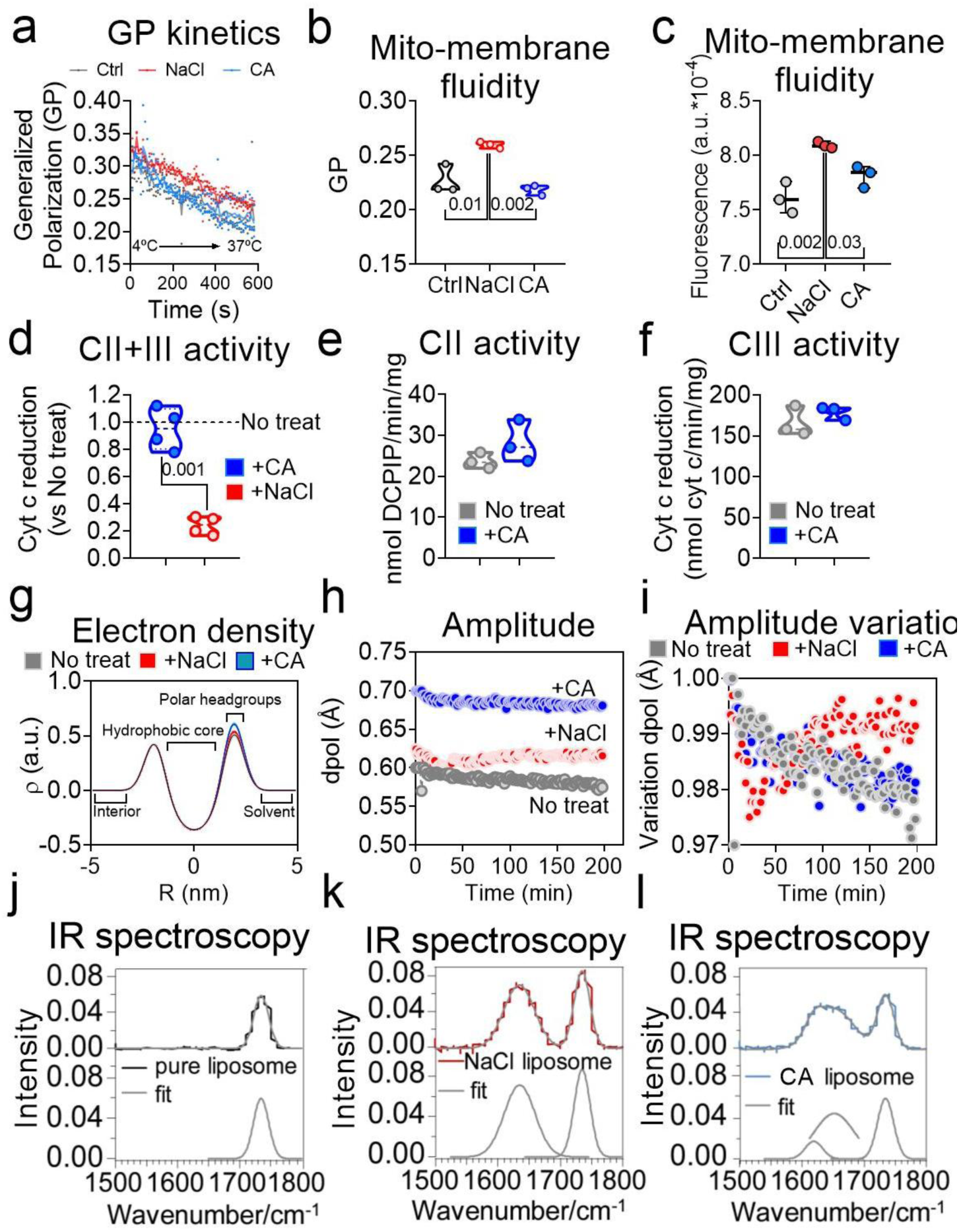
Differential CA interaction with PC permits normal mitochondrial membrane fluidity. **(a)** Fluorescence traces of isolated mitochondrial membranes that were treated without (Ctrl) or with 10 mM NaCl or CA, measured with Laurdan. Temperature was used as internal control (n=3). **(b)** Representation of the minimal values in (a). **(c)** Fluidity quantification of isolated mitochondrial membranes that were treated without (Control) or with 10 mM NaCl or CA, in which MC540 had been incorporated (n=3). **(d)** AA sensitive succinate-driven cyt c reduction of mitochondrial membranes either untreated or treated with 10 mM NaCl or CA. No treat mean was added as a dotted line. **(e)** Succinate-driven DCPIP reduction of mitochondrial membranes with and without 10 mM CA. **(f)** AA sensitive succinate-driven cyt c reduction of mitochondrial membranes or treated with and without CA. **(g)** Difference in electron density profiles for ex-situ samples in either untreated liposomes or treated with 10 mM NaCl or CA, measured by Single-Angle X ray Scattering (SAXS). Labels indicate liposome regions and solvent. **(h)** Final amplitude of the polar electron density layer of untreated liposomes or treated with NaCl or CA measured by SAXS. **(i)** Percentage of amplitude variation over time of untreated liposomes or treated with NaCl or CA measured by SAXS. **(j)** IR absorption spectra of the carbonyl group of PC without or with 10 mM NaCl and CA. Data are presented as mean ± percentiles in (b and d-f) and median ± s.d. in (c) and mean ± percentiles in (c-f). One-way ANOVA analysis was performed in (b and c) and unpaired two-tailed Student’s t-test was used in (d-f).

Given this apparent contradiction, we asked whether CA could interact with lipid bilayers or this behaviour was exclusive for Na^+^. To answer this question, we performed small-angle X-ray scattering (SAXS) of either untreated liposomes, liposomes treated with NaCl or with CA. Overall, the different conditions both in the *ex-situ* prepared samples and in the *in-situ* kinetic measurements were similar, with a slightly stronger interlayer interference produced by Na^+^ (Extended Data Fig. 4a and b). These findings indicated that the liposomes were comparable in size across all conditions, although their degree of multilamellarity varied moderately. Interestingly, the profiles of the liposome’s bilayers treated with NaCl or CA exhibited higher electron-dense polar regions (Fig. 2g), indicating that both Na^+^ and CA were able to not only interact, but also to exclusively rearrange the hydrophilic surface of the bilayer, without altering membrane packing. The extent of the interaction and structural rearrangement of the bilayer surface region was greater for CA than for Na^+^, as evidenced by its higher peak in the electron density plots (Fig. 2g and Extended Data Fig. 4c-d). A time-dependent analysis of the bilayer surface region showed that the amplitude of the polar layer peak was substantially higher for CA than for Na^+^ (Fig. 2h-i), further supporting a stronger interaction and rearrangement of the membrane surface with the small and highly polarizing CA cation. These results indicate that Na^+^ and CA are both able to interact with and exclusively rearrange the polar region of phospholipids in the membrane surface, being CA the strongest interactor. However, in contrast to Na^+^, CA fails to reduce mitochondrial membrane fluidity and electron transfer.

We aimed to understand why despite interacting with phospholipids, CA did not reproduce the physiological effects observed with Na^+^. We reasoned that, given the higher charge density of CA, it was possible that its interaction with phospholipids involved a different functional group in the membrane surface from that engaged by Na^+^. Na^+^ coordinates with the carbonyl groups of phosphatidylcholine (PC) through a lone electron pair located in an sp²-hybridized orbital of their O atoms, forming 1Na^+^:3PC adducts^12,13^. Another negatively charged moiety in PC is the phosphate group, which, given the high electropositivity of CA, represented a potential interaction site for this cation. To measure CA:PC interaction at the intramolecular level we performed infrared (IR) spectroscopy of untreated liposomes, liposomes treated with NaCl or with CA. We confirmed that NaCl-treated liposomes showed the typical left shift in the peak corresponding to the carbonyl group in comparison to untreated liposomes (Fig. 2j and k and Extended Data Fig. 5a and b). However, CA-treated liposomes not only failed to show a shift in the phosphate moiety peak (Extended Data Fig. 5c) but also presented a similar left shift in the carbonyl group region of the IR spectra (Fig. 2c and Extended Data Fig. 5c). Peak deconvolution revealed that this shift comprised two overlapping signals, the major one was closer to the original carbonyl peak than the corresponding signal observed with Na^+^. The appearance of the second peak indicates that CA is able to stablish stronger and shorter bonds with the shared electron pair in the oxygen of the carbonyl group, probably due to its lower atomic radius and higher electropositivity. These series of experiments not only revealed that CA and Na^+^ interact at the same surface level within phospholipids but also suggested that, given the distinct bonds, CA may adopt a distinct coordination geometry with PC relative to Na^+^. Thus, we hypothesised that the lower radius of CA would enable a distinct PC interaction stoichiometry. To measure this, we performed Inductively Coupled Plasma Mass Spectrometry (ICP-MS) of liposomes treated with CA. We found that, in contrast to the 1Na^+^:3PC (Extended Data Fig. 6a)^12,13^, CA interacted in a proportion of 0.21 ± 0.02 atoms per PC molecule, implying that every CA ion coordinates with 5 phospholipids simultaneously (Extended Data Fig. 6b). This disparity in coordination between both cations translated into distinct interaction geometries, resulting in PC super-assemblies with different structural arrangements (Extended Data Fig. 6c and d). Altogether, these data indicate that, whereas Na^+^ forms an edged, trigonal adduct with PC that hinders phospholipid movement across the IMM, CA forms a more stable penta-coordinate structure, which allows unimpeded mitochondrial membrane fluidity, ultimately enabling mitochondrial electron transfer.

Given its stronger interaction with phospholipids, we hypothesised that CA may replace mitochondrial Na^+^ in PC coordination under relevant (patho)physiological conditions, imposing its characteristic penta-coordinate membrane surface geometry and restoring the physiological parameters associated with hindered mitochondrial electron transfer. We measured whether CA, applied in clinically relevant concentrations, could reach the mitochondrial matrix. ICP-MS showed that that this cation accumulated 2.58 times more in the mitochondria from cells incubated with CA. This co-existed with matrix Na^+^ in the G11778A mutant model (Fig. 3a). Mechanistically, CA’s stronger interaction with phospholipids (Fig. 2g-l and Extended Data Fig. 4 and 5) was predicted to substitute Na^+^ in the IMM (Fig. 3b), while the ability of CA to establish a pentameric membrane surface geometry was sufficient to rescue G11778A IMM fluidity (Fig. 3c and Extended Data Fig. 7a), which translated into an increase of the CII+III electron transfer (Fig. 3d and e). Importantly, these effects were reproduced by adding CA to the reaction mixture containing isolated G11778A mitochondrial membranes, immediately before CII+III activity (Fig. 3f) or fluidity (Fig. 3g) measurements. These results indicate that this response was independent from other possible CA targets, as they were not detected by proteomics analyses of these samples. Unimpeded electron transfer between CII and CIII resulted in the recovery of H^+^ pumping by CIII and CIV and a concomitant reestablishment of ΔΨmt (Fig. 3h). In parallel, promoting penta-coordinate geometry in the IMM by the application of CA decreased ROS production in the G11778A mutant model to the levels of its isogenic control (Fig. 3i). As a consequence, HIF-1α stabilization and activity decreased in G11778A (Extended Data Fig. 7b and c), which was readily evident as media acidification diminished under CA treatment (Extended Data Fig. 7d). In addition, G11778A cell proliferation returned to control levels upon CA treatment (Fig. 3j). All these occurred without altering electron entry into the OxPhos system through CI (Extended Data Fig. 7e). So far, these data show that G11778A is a genetic model showing the constitutive activation of the Na^+^-hindered electron transfer mechanism. Also, imposing a CA-dependent pentameric PC assembly over the trigonal adduct elicited by Na^+^ demonstrates that membrane surface geometry is a decisive factor for respiratory chain activity, ROS production and HIF-1-dependent adaptation in live cells.

**Figure 3.**
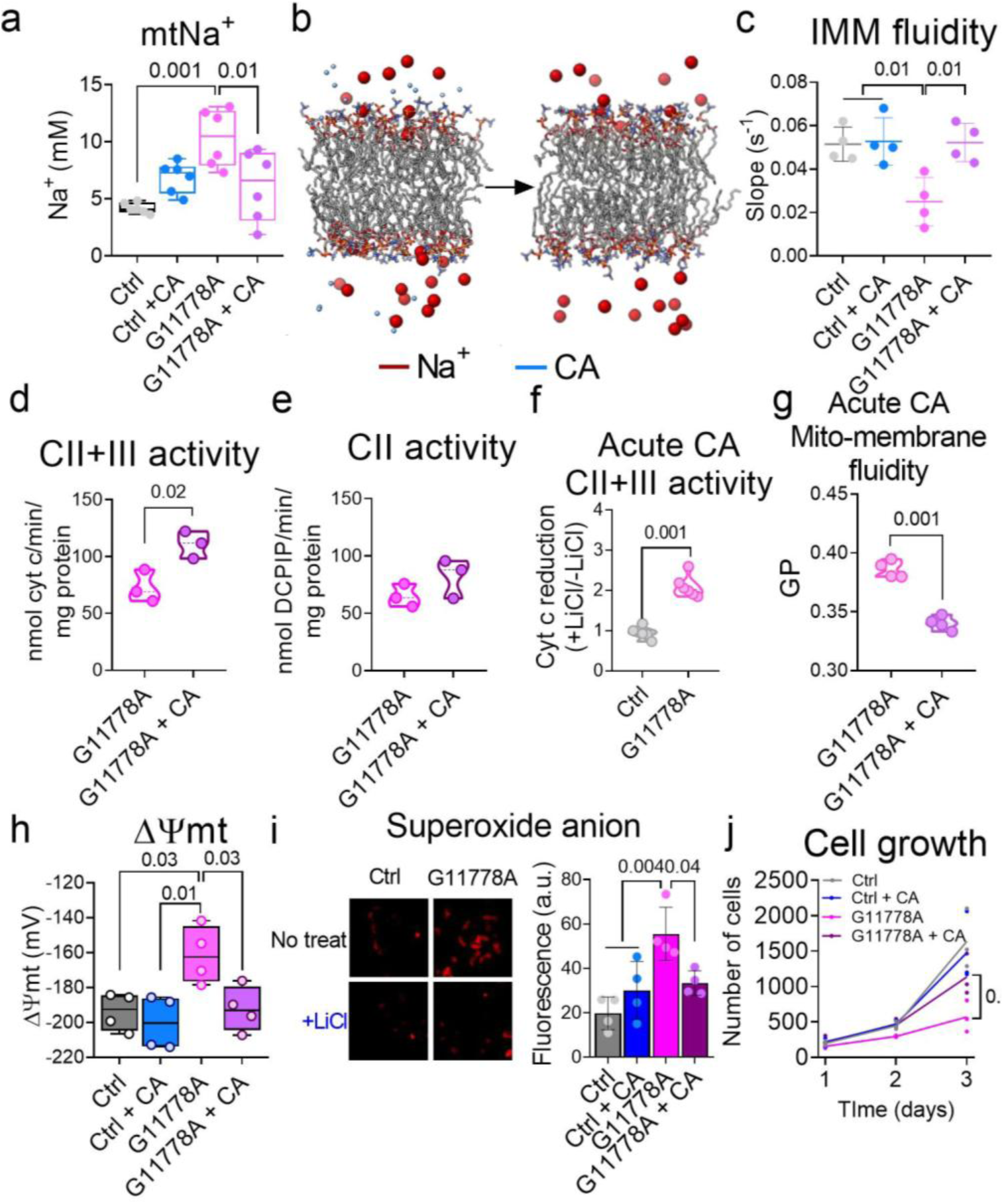
CA rescues electron transfer and the hypoxic signature in G11778A. **(a)** Mitochondrial matrix Na^+^ measured by confocal microscopy in G11778A cybrids and its isogenic control (Ctrl) either non-treated or chronically treated with 1 mM CA. **(b)** The first (left) and last (right) frames from a 200 ns molecular dynamics simulation of a POPC membrane in the presence of Na^+^ and CA. **(c)** Quantification of FRAP of Ctrl and G11778A cells treated with and without 1 mM CA. **(d)** AA sensitive succinate-driven cyt c reduction of mitochondrial membranes from either untreated or 1mM CA-treated control or G11778A cells. **(e)** Succinate-driven DCPIP reduction of mitochondrial membranes from either untreated or 1mM CA-treated control or G11778A cells. **(f)** AA sensitive succinate-driven cyt c reduction of G11778A mitochondrial membranes with or without 10 mM CA in the reaction mixture. **(g)** Fluidity of isolated mitochondrial membranes from G11778A in the absence (Ctrl) or presence of 10 mM CA, measured with Laurdan. **(h)** ΔΨmt measured by confocal microscopy in control and G11778A cells either untreated or treated with 1 mM CA. **(i)** Superoxide production measured by confocal microscopy in control and G11778A cells either untreated or treated with 1 mM CA. Left: representative images of superoxide staining; right: quantification of fluorescence. **(j)** Proliferation of G11778 and Ctrl cells in the absence or presence of 1 mM CA. Data are presented as mean ± s.d. in (a, c, h and i) and mean ± percentiles in (d-g). One-way ANOVA analysis was performed in (a, c, h and i) and unpaired two-tailed Student’s t-test was used in (d-g).

Finally, we asked whether imposing a penta-coordinate membrane surface geometry would interfere with redox signalling in acute hypoxia, a physiological adaptive mechanism in which the Na^+^:phospholipid interaction is essential^12^. For that, we added CA immediately before hypoxic incubation and several parameters associated with hypoxic adaptation were subsequently assessed. Acute CA addition on cells prevented the decrease in CII+III activity associated with the hypoxic formation of Na^+^-dependent trigonal adducts (Fig. 4a), as well as the reduction in IMM fluidity (Extended Data Fig. 7g and Fig. 4b), which translated into the prevention of hypoxic mitochondrial depolarization (Fig. 4c and d). In summary, these results demonstrate that the modulation of IMM surface geometry, through the alteration of the mitochondrial cation composition, is able to regulate mitochondrial energy conversion, hypoxic redox signalling and adaptation.

**Figure 4.**
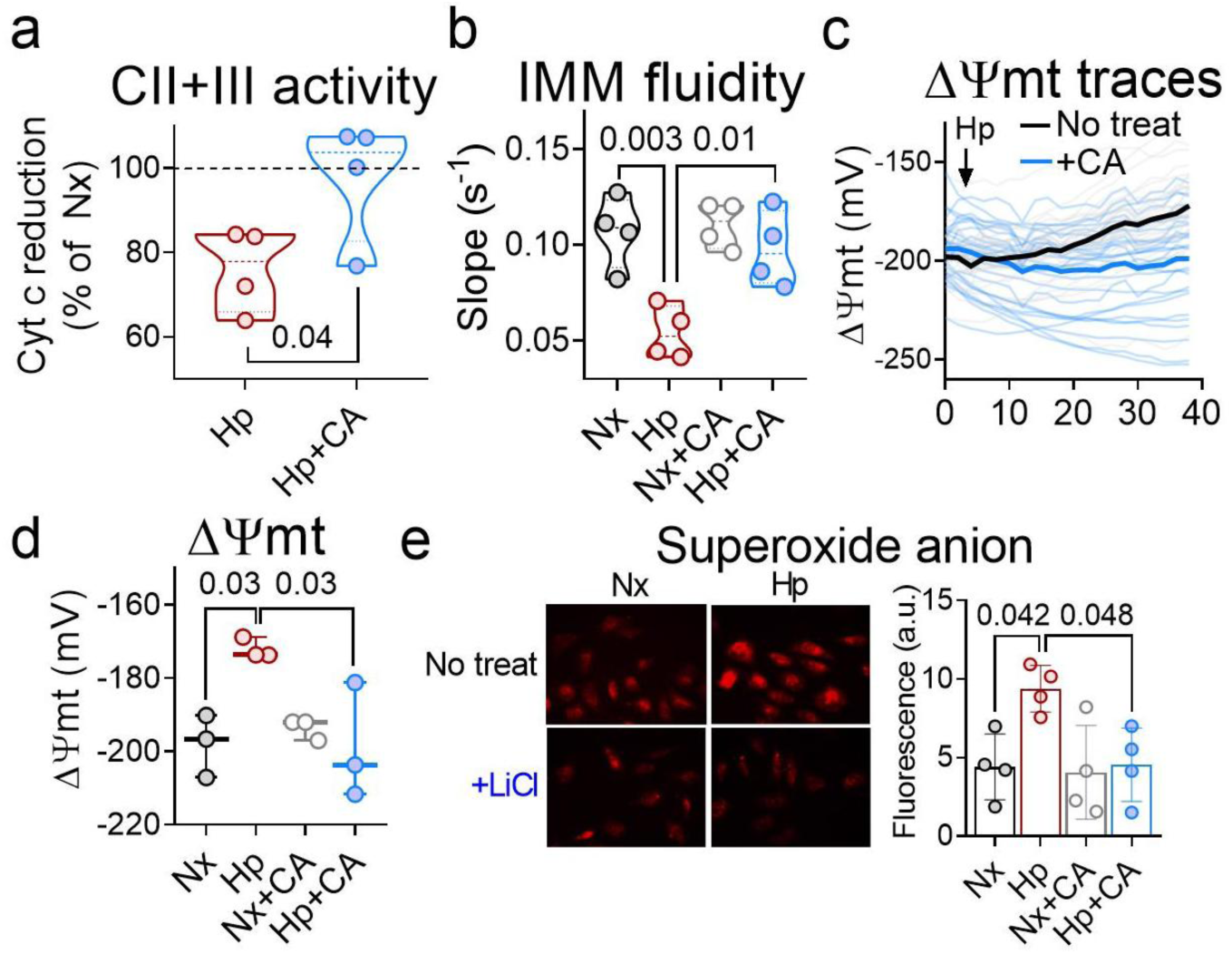
Acute CA treatment restores mitochondrial electron transfer and hypoxic redox signalling. **(a)** AA sensitive succinate-driven cyt c reduction of mitochondrial membranes from either untreated or 1mM CA-treated bovine aortic endothelial cells (BAECs) that had been subjected to 15 minutes of hypoxia (Hp; 1% O_2_). **(b)** Quantification of FRAP of BAECs treated with and without 1 mM CA that had been subjected to 15 min of normoxia (Nx) or Hp. **(c)** ΔΨmt traces of BAECs either untreated (Ctrl) or acutely treated with 1 mM CA that were subjected to Hp, measured by confocal microscopy (n=3; grey: individual Ctrl cells; light blue: individual CA-treated cells). **(d)** ΔΨmt values of BAECs either untreated (Ctrl) or acutely treated with 1 mM CA that were subjected to 30 min of Nx or Hp, measured by confocal. **(e)** Superoxide detection after incubation with dihydroethidium (DHE) in normoxic or hypoxic 10-min time windows in non-treated (No treat) or CA-treated (+CA) BAECs. Data are presented as mean ± percentiles in (a and b) and median ± s.d. in (d). One-way ANOVA analysis was performed in (b, d and e) and unpaired two-tailed Student’s t-test was used in (a).

The behaviour of ions in biological solutions, particularly regarding to their interactions with bio-membranes, remains a largely unexplored area of research. Phospholipid bilayers are embedded in solutions containing multiple dissolved ions. Some efforts have been made to understand how ions affect membrane homeostasis at the atomic or sub-molecular level^13–17^. These approximations have predicted that ion:phospholipid interaction vary depending on each chemical species. In this way, Na^+^ was predicted to interact with PC at its carbonyl group^13^, which we confirmed to occur and was observed to have a relevant role in physiology and disease^11,12,18–22^. Several studies have reported a stronger interaction of CA with PC^23,24^, which basis was proposed to rely on the shorter ionic radius distance between the shared electron pair and the highly electropositive CA. From these studies, it might be expected that, given that CA forms more stable bonds, its impact on membranes would surpass that of Na⁺, further reducing IMM fluidity, decreasing electron transfer and increasing mtROS production. However, while we confirmed that CA interacts more strongly with the carbonyl group of PC than Na⁺, IMM fluidity and CoQ transfer remained unaffected in its presence in physiological conditions. This apparent contradiction was resolved by observing that CA forms a penta-coordinate multi-PC assembly that does not impede the lateral diffusion of other phospholipids/lipids within the membrane, whereas the tri-coordinate PC adduct, specifically formed by Na⁺, restricts mobility in the IMM (Extended Data Fig. 8). Such behaviour is explained by the fact that the CA:PC pentamer possesses, from the top-down view, five axes of symmetry (at 72° intervals), in contrast to the three axes of the trigonal Na^+^:PC adduct (120°). This higher degree of isotropy promotes favourable orientations for lateral diffusion in multiple directions. This previously unrecognized property of membrane biology enables pentagonal surface geometry, compared to a trigonal assembly, to sustain IMM fluidity and CoQ transfer, restraining the production of mtROS and the activation of the hypoxic redox signalling cascade.

In summary, these data show that multi-phospholipid assemblies are modulated by the composition of the surrounding solution, whereby cations are capable of imposing specific coordinates with IMM phospholipids to form a variety of multi-PC structures regulating energy conversion, mtROS production and adaptive responses. In this way, membrane surface geometry emerges as a critical and an unexpected determinant of membrane function, governing mitochondrial biology, with broad implications for physiology and disease.

## Methods

### Cell culture and transfection

Cells were routinely maintained in cell culture incubators (95% air, 5% CO_2_ in gas phase, at 37 °C). BAECs were isolated as previously described^12^ and cultured in RPMI 1640 supplemented with 15% heat-inactivated FBS, 100 U/mL penicillin and 100 μg/mL streptomycin. Slc8b1 (NCLX)fl/fl (NCLX^WT^) and Slc8b1 (NCLX)-/-(NCLX^KO^) mouse embryonic fibroblasts (MEFs) were isolated and immortalized as described previously^12,25^. Ctrl and G11778A cybrids were kindly donated by Dr. José Antonio Enríquez. NCLX^WT^, NCLX^KO^, Ctrl and G11778A were cultured in DMEM supplemented with 10% heat-inactivated FBS, 100 U/mL penicillin and 100 μg/mL streptomycin. Human umbilical vein endothelial cells (HUVECs) were isolated as previously described^12^ and cultured in Medium 199 supplemented with 20% heat-inactivated FBS, 16 U/mL heparin, 100 mg/L endothelial cell growth factor (ECGF), 20 mM HEPES, 100 U/mL penicillin and 100 μg/mL streptomycin. BAECs were used between passages three and nine, and HUVECs between passages three and seven. For MEFs were correctly identified visually and endothelial morphology was assessed by visual inspection, and cross-contamination/invasion with fibroblasts or other cell types was tested negative by checking endothelial nitric oxide synthase (eNOS) expression. 1 mM CA treatment was performed chronically unless otherwise stated.

Cells were transfected with pDsRed2-Mito vector (Clontech) as 0.25 μg of vector DNA per 0.8 cm^2^ well was carried out using Lipofectamine 2000 (Invitrogen).

### Detection of superoxide by fluorescence microscopy in fixed cells

Cells were plated on glass coverslips one day prior to the experiments. In hypoxic experiments, 30 mM CA was introduced 30 minutes before the start of the experiment and remained present throughout the experimental procedure. In experiments involving G11778, 1 mM CA was applied chronically and remained present throughout the experimental procedure. For hypoxia treatments, all solutions were pre-equilibrated under hypoxic conditions before application. The plated cells were then transferred into an Invivo2 400 workstation (Ruskinn) maintained at 1% O₂, 5% CO₂, and 37°C, where they were incubated for specified durations (0, 15, 30, 45, and 60 minutes) in fresh medium. Following incubation, the cells were washed three times with Hank’s balanced salt solution containing Ca²⁺/Mg²⁺ (HBSS + Ca/Mg) and then incubated with 5 μM dihydroethidium (DHE) in HBSS + Ca/Mg for 10 minutes in the dark. After removing excess probe by three additional washes with HBSS + Ca/Mg, the cells were fixed with 4% paraformaldehyde (PFA) and kept in the dark at 4°C for 15 minutes. Post-fixation, the cells were washed three more times with HBSS + Ca/Mg, and the coverslips were mounted onto slides. For normoxic treatments, the medium was replaced with fresh normoxic medium, and cells underwent the same procedures as the hypoxic group but were kept in a standard cell incubator. 10 μM Antimycin A was added 30 min prior the start of the experiment and was maintained through as a positive control.

Images were captured using a Leica DMR fluorescence microscope equipped with a 63x objective and 546-12/560 nm excitation/emission filter pairs. Three images were taken per coverslip, with the number of independent experiments detailed in the figure. Quantification was performed using ImageJ software (NIH), applying the same threshold to all images. The mean values from histograms were averaged across the three images obtained from each coverslip.

### Detection of intramitochondrial Na+ and ΔΨmt by live-imaging confocal microscopy

Cells were seeded in eight-well plates one day before experimentation, after which the plated cells were washed three times with HBSS + Ca/Mg with glucose, in the presence or absence of 1 mM CA, and incubated with either 1 μM RedNa Chloride or 30 nM tetramethylrhodamine methyl ester (TMRM) for 30 min at 37°C in the dark. Following incubation, RedNa Chloride was washed out and replaced with fresh HBSS + Ca/Mg. Subsequently, the plate was transferred to a Leica SP8 inverted microscope, an automated stage for live imaging, and a thermostated hypoxic chamber. After focusing the imaging planes, live imaging was acquired every 2 min over 40 min (21 total cycles) using a 20x objective. Normoxia experiments were conducted at constant 20% O₂ and 5% CO₂, while hypoxia experiments began at 20% O₂ and 5% CO₂ before transitioning to 2% O₂ and 5% CO₂ at the second cycle. RedNa chloride fluorescence was detected using the 514 nm line and acquired using the 555–575-nm range, while TMRM was excited/acquired at 545/555-590 nm respectively.

Individual cell quantification was performed using Fiji software. For each experiment and condition, all cells in the plane were classified as regions of interest (ROIs) and quantified, the mean fluorescence per cell was recorded for each time cycle.

### Western blot analysis

Protein samples were extracted using non-reducing Laemmli buffer (without bromophenol blue) and quantified via Bradford assay. Following quantification, the extracts were mixed with 5% 2-mercaptoethanol and loaded onto 10% standard polyacrylamide gels for electrophoresis, after which the separated proteins were transferred to either nitrocellulose or PVDF membranes. The membranes were probed with the following primary antibodies: mouse monoclonal anti–HIF–1α, kindly provided by Dr. Silvia Martín Puig (IIB-SM), mouse monoclonal NDUFA4L2 (NUOMS; clone 1G1H10; 66050; Proteintech) and mouse monoclonal anti-β-actin antibody (clone AC-15; A3854, Sigma). Antibody binding was detected using species-specific fluorescein-conjugated secondary antibodies followed by fluorescence visualization on a Odissey digital luminescent image analyser.

### Measurement of oxygen consumption

The oxygen consumption rate (OCR) was assessed using an XF96 Extracellular Flux Analyzer (Seahorse Bioscience), with Ctrl and G11778A plated at 6 × 10^2^ cells per well one day prior to experimentation. Cells were preincubated for 1 hour at 37°C in unbuffered DMEM containing 25 mM glucose, 1 mM pyruvate, and 2 mM glutamine in a CO₂-unregulated incubator. The OCR measurement protocol involved sequential injections of: unbuffered DMEM (baseline), 5 μg/mL oligomycin (ATP synthase inhibitor), 300 nM FCCP (mitochondrial uncoupler), and 1 μM rotenone plus 1 μM antimycin A. OCR values were normalized to percentage of living cells determined by CyQuant assay. Basal respiration was measured first, followed by oligomycin treatment to determine ATP-linked respiration (calculated as basal OCR minus oligomycin-inhibited OCR, representing coupling efficiency). FCCP-induced uncoupling revealed maximal respiratory capacity (reflecting mitochondrial reserve capacity), while subsequent rotenone/antimycin A treatment quantified non-mitochondrial oxygen consumption (remaining OCR after complete ETC inhibition). All calculations followed manufacturer protocols, with oligomycin-sensitive respiration indicating OXPHOS activity, FCCP response showing ETC capacity, and rotenone/antimycin-resistant respiration representing non-mitochondrial oxygen consumption.

### Cell proliferation

Cell proliferation was assessed by manual cell counting. Ctrl and G11778A cells were seeded at a density of 200 cells per well in 48-well plates or 800 cells per well in 6-well plates. At the indicated time points, one well per experimental condition was trypsinized each day, and the resulting cell suspension was collected. Cell numbers were subsequently determined manually using a Neubauer haemocytometer. Proliferation curves were generated by plotting the number of cells counted at each time point.

### Mitochondria isolation

Mitochondria were isolated from Ctrl, G11778A and MEFs using a cell culture-adapted protocol^1^, which involved resuspending a fresh cell pellet in sucrose buffer within a glass Elvehjem potter, followed by homogenization through multiple up-and-down strokes using a motor-driven Teflon pestle; subsequent sequential homogenization and centrifugation steps were then performed to obtain the mitochondrial fraction.

### Mitochondrial membrane isolation and complex activity measurement

Mitochondrial membranes were prepared from Ctrl, G11778A or BAECs by subjecting isolated mitochondria to freeze-thaw cycles, followed by measurement of OxPhos enzyme activities according to established methods^1^

Using approximately 20 μg of protein per sample, CII (succinate dehydrogenase) activity was determined in a reaction mixture containing mitochondrial membranes, succinate, and DCPIP by tracking absorbance changes at 600 nm. CII+III activity was quantified as antimycin A-sensitive succinate-cytochrome c reduction at 550 nm. Isolated CIII activity was measured as antimycin A-sensitive ubiquinone 2-cytochrome c reduction at 550 nm. Isolated CI activity was measured as rotenone-sensitive NADH-decylubiquinone reduction at 340 nm. All assays were performed in the presence or absence of 10 mM CA or 10 mM NaCl.

### Membrane fluidity assessment in isolated mitochondrial membranes

Mitochondrial membrane fluidity from Ctrl and G11778A samples, either untreated or treated with 10 mM NaCl or 10 mM CA was assessed using the ratiometric fluorescent probe Laurdan (6-dodecanoil-2-dimethylaminonaftalene) or Merocyanine 540 (MC540). The probes were first dissolved in DMSO at a concentration of 1mM as a storage stock solution. A working stock was then prepared by adding a final concentration of 10 μM of the storage stock to Medium A (0,32 M sucrose, 10 mM Tris-HCl, 1 mM EDTA, pH=7,2) supplemented with 3 mg/ml BSA. In 96-well plates, 20 μg mitochondrial membrane samples were incubated with 100 μl of the working stock for an hour at room temperature in the dark. After incubation, samples were centrifuged at 16.000 rcf for a minute at RT, supernatant was discarded and the pellet was washed with Medium A with 3 mg/ml BSA. The centrifugation and washing were repeated three times. Finally, the samples were resuspended in Medium A with BSA using a vortex and the 96-well plate was kept at 4°C for 10 minutes before measuring fluorescence at a CLARIOstar Plus Microplate Reader.

Using a 350 nm excitation wavelength, emission was collected at 440 nm and 490 nm wavelengths during 50 cycles (intervals of 4 seconds) for Laurdan. For MC540, samples were excited at 550 nm and emission was collected at the interval 560-580 nm during 50 cycles. The microplate reader’s thermostat was turned on just before the first measurement, allowing the sample to increase its temperature gradually from 4°C to 37°C. Fluorescence values were obtained as an average of technical duplicates. These values were then analysed by calculating the Generalised Polarization or GP using the equation GP= (I440-I490)/(I440+I490); where I440 and I490 represent the fluorescence at 440 and 490 nm respectively.

### Fluorescence quenching recovery after photobleaching (FRAP)

Ctrl, G11778 or BAECs, either untreated or treated with 1 mM CA, were transfected with the pDsRed2-Mito vector (Clontech), after which the growth medium was replaced with HBSS + Ca/Mg + glucose, with or without 1 mM CA, and the plate was transferred to a Leica SP-5 confocal microscope equipped with an automated stage for live imaging and a thermostated hypoxic chamber. Imaging planes were focused, and images were acquired using a 63x objective with 13x zoom. Samples were excited using the 514-nm line of an argon/krypton laser, with emission detected between 565–595 nm, and images were captured using Leica TCS software. For FRAP analysis, MitoRFP was scanned five times before photobleaching was performed with 15 scans at 40% laser power. Fluorescence recovery was then monitored over 60 sequential scans acquired at 1-second intervals. FRAP measurements under normoxic conditions were conducted at 20% O₂ and 5% CO₂, after which the chamber was switched to 1% O₂ and 5% CO₂, and a second FRAP measurement was performed following a 15-minute hypoxic equilibration period.

### Fourier-transform IR spectroscopy

Phosphatidylcholine (PC) liposomes were prepared via the thin-film hydration method, followed by sequential extrusion through polycarbonate filters of decreasing pore sizes (400 nm, 200 nm, and 100 nm) as described in reference 2. The lipid film was hydrated using high-purity water with minimal metal impurities (Optima LC/MS Grade, Fisher Chemical), and the resulting liposomes were concentrated by filtration to achieve a final lipid concentration of 72 mg/mL, as quantified by the Rouser assay46. For metal binding studies, CA and sodium salts were mixed with the liposomes at a molar ratio of 16 mM cation:0.5 mg/mL lipids and incubated at 37°C for 2 hours. Fourier-transform infrared (FTIR) spectroscopy was subsequently performed using a Nicolet 6700 spectrometer (Thermo Scientific) in transmission mode to analyse both plain liposomes and metal-incubated liposomes in liquid phase. All experimental procedures were conducted in triplicate to ensure reproducibility.

### Inductively coupled plasma mass spectroscopy

PC liposomes were incubated with sodium chloride at 37°C for 2 hours at a stoichiometric ratio of 16 mM cation:0.5 mg/mL lipids. Mitochondria from untreated or 1 mM CA-treated NCLX^WT^ or NCLX^KO^ MEFs were isolated and the mitochondrial pellet was stored. 18 p150 culture dishes were collected per condition. Following incubation, all samples underwent three sequential washing steps via filtration using Optima LC/MS Grade water (Fisher Chemicals) to remove unbound ions, followed by acid digestion with ultra-pure HNO3 (13 ppt CA content, Optima grade for trace metal analysis, Fisher Chemicals). The quantity of CA cations bound per lipid molecule was quantified using a Thermo Scientific iCAP-Q inductively coupled plasma mass spectrometer (ICP-MS) equipped with collision/reaction cell technology and kinetic energy discrimination capability. Concurrently, lipid concentrations were verified through the Rouser assay^26^. All experimental procedures were performed in triplicate to ensure reproducibility.

### Small Angle X-ray scattering (SAXS)

The experiments were carried out at the BL11 NCD-SWEET beamline, which is a dedicated Small-Angle X-ray scattering (SAXS) beamline at the ALBA Synchrotron in Barcelona, Spain, under project number 20260380105. Sample holders were borosilicate capillaries with a wall thickness of ten micrometres and an external diameter of two millimetres, and these capillaries were placed in a temperature-controlled holder maintained at 37 degrees Celsius. To monitor how the samples evolved over time, 5 seconds SAXS patterns were collected every two minutes during 2 hours after mixing.

For the modelling of the electron density profile, the liposomes are treated as locally planar because their size is much larger than the bilayer thickness. The profile itself is built asymmetrically from the sum of three Gaussian functions, with two positive Gaussians representing the outer and inner polar headgroup regions and one negative Gaussian representing the hydrophobic acyl-chain core^27^. The key distinction here is that the amplitude of the outer polar region is allowed to differ from that of the inner one, which explicitly accounts for any asymmetry between the two sides of the membrane^28^. The scattering intensity for a single bilayer is then obtained by taking the square of the Fourier transform of this profile in Cartesian coordinates. However, because the samples exhibited some degree of multilamellarity, a further difference in the modelling is introduced: a small fraction of the total scattering is instead treated as multilamellar stacks, for which a Caille structure factor is used to incorporate the effects of bilayer fluctuations and interlayer correlations^29,30^. This separate treatment distinguishes the strictly single-bilayer contribution from the more complex, stacked arrangement that is also present in the samples.

*In situ* kinetics correspond to the experimental setup where the liposome suspension and saline solutions were admixed at the moment of analysis. *Ex situ* samples, however, denote those that had already been prepared beforehand and kept refrigerated prior analysis. This difference reflects the consistency of cation:membrane interaction during time.

### Proteomic analysis

Protein extracts of whole cells or isolated mitochondria from Ctrl and G11778A cells were obtained by homogenization with ceramic beads (MagNa Lyser Green Beads, Roche, Germany) in CS buffer (Pipes pH6.8, MgCl2, NaCl, EDTA, sucrose, SDS, sodium orthovanadate; Biochain Institute, Inc. #K3013010-5) freshly supplemented with protease and phosphatase inhibitors. Extracted proteins (around 200 μg) were subjected to in-filter reduction and alkylation using iodoacetamide followed by trypsin digestion (Nanosep Centrifugal Devices with Omega Membrane-10K, PALL), and the resulting peptides were TMT-labeled according to the manufacturer’s instructions. Labeled peptides were loaded and washed on Evotips for chromatographic separation by an evosep one HPLC system (30 SPD method, with Endurance Column 15 cm x 150 μm ID, 1.9 μm beads-EV1106, Evosep). Mass spectra were acquired in a data-dependent manner, with an automatic switch between MS and MS/MS using a top-speed method and dynamic exclusion. MS spectra were collected in the Orbitrap analyser using a mass range of 375–1500 m/z at 60,000 resolution. HCD fragmentation was performed at 33 eV of normalized collision energy and MS/MS spectra were analysed at 30,000 resolution in the Orbitrap.

Proteins were identified with the SEQUEST HT algorithm integrated in Proteome Discoverer 2.5 (Thermo Scientific). MS/MS scans were searched against a pig reference target-decoy protein database (human_pig_202105_pro-sw-tr.target-decoy.fasta), (296316 sequences in total). For database searching, parameters were selected as follows: trypsin digestion with 2 maximum missed cleavage sites, precursor mass tolerance of 2 Da, and a fragment mass tolerance of 0.03 Da. Methionine oxidation (+15.994915 Da) and asparagine and glutamine deamidation (+0.984016 Da) were set as variable modifications, whereas cysteine carbamidomethylation (+57.021464 Da) and TMT labeling (+229.162932 Da) at peptide N-terminal ends and Lys residues were considered fixed modifications. False discovery rates (FDR) for peptide identifications were calculated by the refined method43 after additional filtering for a precursor mass tolerance of 10 ppm.44 A 1% FDR was used as criterion for peptide identification. Proteins showing differential expression between WT and KO were identified using limma v-3.50.3 (refence 1) based on Zq values.

Quantitative information from TMT reporter intensities was integrated from the spectrum level to the peptide level and then to the protein level based on the WSPP model,45,46 using the GIA integration algorithm.47

To note, taking into account that the protein extraction protocol is not specific for membrane proteins, the peptides of these proteins are poorly represented in this study. This includes ND6 or ND4 in all models analyzed.

Protein abundance analysis

Differential protein abundance was assessed with the limma framework^31^, fitting a linear model in which genotype served as the factor of interest. Proteins with an adjusted p-value (Benjamini–Hochberg correction) of padj ≤ 0.05 were considered statistically significant. As before, gene annotation was performed using the Bioconductor R packages biomaRt (v2.58.2) and org.Mm.eg.db (v3.22.0).

Gene Set Enrichment Analysis (GSEA)^32^ was applied to identify Gene Ontology biological process signatures that were differentially regulated across experimental conditions. Enrichment analysis was carried out using the clusterProfiler package (v4.10.1)^33^, with false discovery rates controlled using the Benjamini–Hochberg correction.

### Molecular dynamics simulations

The initial membrane system was generated using the Membrane Builder module of CHARMM-GUI^34^. The system consisted of a POPC lipid bilayer containing 30 lipid molecules per leaflet, parameterized using the CHARMM36m force field. The membrane was placed in a cubic simulation box with a minimum distance of 1.0 nm between the solute and the box boundaries and solvated with TIP3P water molecules. CA and NaCl were added to a final concentration of 0.15 M each. Molecular dynamics simulations were performed using GROMACS 2025.3 (https://doi.org/10.1016/j.softx.2015.06.001)^35^. Energy minimization was carried out using the steepest descent algorithm. The system was equilibrated at 310 K in two consecutive NVT stages (250 ps total) using the V-rescale thermostat, followed by three NPT equilibration stages under semi-isotropic pressure coupling using the C-rescale barostato. The first NPT stage was performed for 125 ps with a 1 fs integration timestep, followed by two additional stages of 500 ps each using a 2 fs timestep. Position restraints were gradually reduced throughout the equilibration protocol. Long-range electrostatic interactions were treated using the Particle Mesh Ewald (PME) method with a real-space cutoff of 1.2 nm. A cutoff of 1.2 nm was also applied for van der Waals interactions. Covalent bonds involving hydrogen atoms were constrained using the LINCS algorithm. Production simulations were performed for 200 ns using a 2 fs integration timestep under periodic boundary conditions. Unless otherwise stated, all remaining simulation parameters were those defined by the default CHARMM-GUI membrane simulation protocol. Cation:phospholipid interaction was visualized using Winmostar V11.16.3.

### Statistics

The results are expressed as mean ± standard deviation of the mean (s.d.) unless otherwise indicated. Prior to statistical analysis, all datasets were assessed for normality (Shapiro-Wilk or Kolmogorov-Smirnov tests) and homogeneity of variance (Bartlett’s or Levene’s tests). For multi-group comparisons, one-way analysis of variance (ANOVA) was performed followed by either Tukey’s honest significant difference (HSD) or Bonferroni post-hoc tests to evaluate all experimental groups, unless otherwise stated. Two-group comparisons were conducted using Student’s two-tailed t-test for normally distributed data. Statistical significance was defined as p < 0.05 for all analyses, with exact p-values reported in the figures when significant. All statistical analyses were performed using GraphPad Prism version 8 and 11 (GraphPad Software) statistical package.

### Reporting summary

Further information on research design is available in the Nature Research Reporting Summary linked to this paper.

## Acknowledgements

We thank L. Fernandez-Mendez (CIC biomaGUNE) for support in liposome synthesis; C. Rueda and J. Satrústegui (CMBSO, UAM-CSIC), L. del Peso (UAM) and P. Bernardi (UP) for helpful discussions. We thank the Microscopy unit service and the Molecular and Cellular Biology (CNC) for their support.

Project carried out with a 2024 Leonardo Grant for Scientific Research and Cultural Creation from the BBVA foundation (LEO25-1-17476-BBM-BAS-2) and supported by a RYC2022-036516-I (MICIU) to P.H.-A, FPU18/03475 to C.C.-F. (MICIU). J.R.-C. was supported by PID2024-155807OB-C21. SCR was supported by PID2022-142842OB-I00 and RYC2020-030241-I. Scheme figures were made with BioRender.

## Contributions

P.H.-A. and C.C.-F. designed the study. C.C.-F., M.G.-H., D.C.-R. and P.H.-A. performed the bulk of the experiments and analysed the data. E. C. and J. V. performed the proteomics. J.L.C.-A. analysed the proteomic data. S.C.-R. and J.R.-C performed IR experiments and ICP mass spectrometry experiments. R.A.-P. performed the respiration assays. J.L. performed IR spectroscopy experiments. C. H.-I. performed SAXS experiments. J.W.E. helped with crucial experimental procedures and analysis of the data. P.H.-A and J.W.E. supervised the study. C.C.-F. and P.H.-A. wrote the manuscript.

