## Supplementary figures for "Membrane surface geometry is a determinant of mitochondrial electron transfer and cellular adaptation"

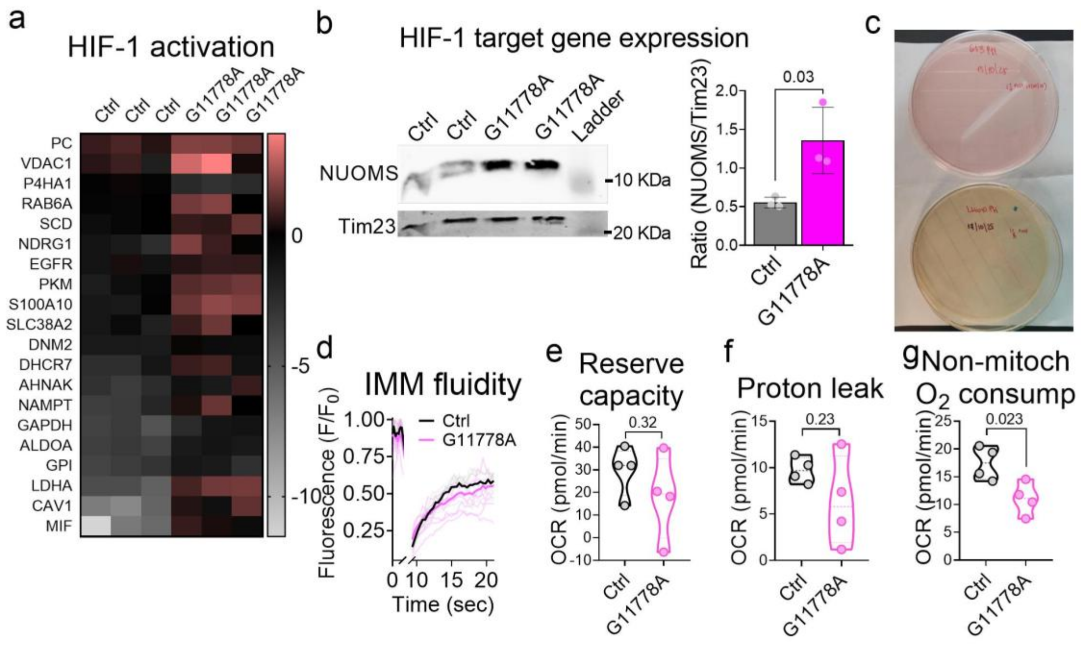


**Extended Data Figure 1. G11778 mutant shows basally activated hypoxic response**. **(a)** Unbiased proteomics analysis of G11778A and its isogenic control (Ctrl) showing the increased presence of HIF-1 targets in the former. **(b)** SDS-PAGE assessing the HIF-1 target NUOMS and Tim23 levels in G11778A versus its isogenic control (Ctrl). **(c)** Representative image of culture dishes containing G11778A cells (down) and its isogenic control (up) at confluence. **(d)** FRAP traces of G11778A and its isogenic control (Ctrl; n=4 independent experiments). **(e)** Proton leak of G11778A and its isogenic control (Ctrl) measured by Seahorse. **(f)** Non-mitochondrial $O_2$ consumption of G11778A and its isogenic control (Ctrl) measured by Seahorse. **(j)** Maximal respiration of G11778A and its isogenic control (Ctrl) measured by Seahorse. **(g)** Non-mitochondrial $O_2$ consumption of G11778A and its isogenic control (Ctrl) measured by Seahorse. Western blot quantifications are shown in the insets of panels b. Data showing an adjusted p-value >0.05 is shown in (a). Data are presented as mean ± s.d. in (b) and as mean ± percentiles in (e-j). Unpaired two-tailed Student's t-test was used in (b and e-j).

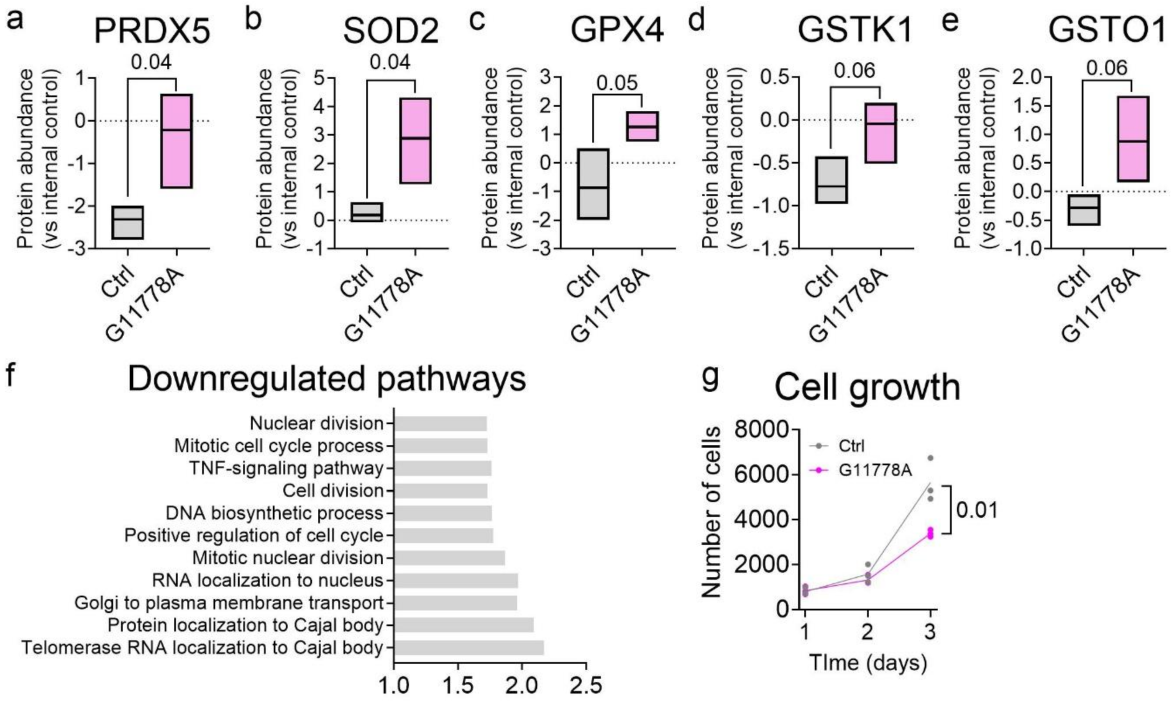


**Extended Data Figure 2. G11778A shows increased expression of antioxidant enzymes and lowered cell proliferation**. **(a-e)** Unbiased proteomics analysis showing the increased presence of (a) peroxiredoxin 5 (PRDX5), (b) superoxide dismutase 2 (SOD2), (c) glutathione peroxidase 4 (GPX4), (d) Glutathione S-transferase kappa 1 (GSTK1) and Glutathione S-transferase omega-1 (GSTO1) in G11778A in comparison to its isogenic control (Ctrl; n=3). **(f)** GSEA showing the downregulated pathways in G11778A in comparison to Ctrl (n=3). **(g)** Proliferation of G11778 and Ctrl cells. Data are presented as median ± percentiles in (a-e) and mean in (g). Unpaired two-tailed Student's t-test was used in (a-e and g).

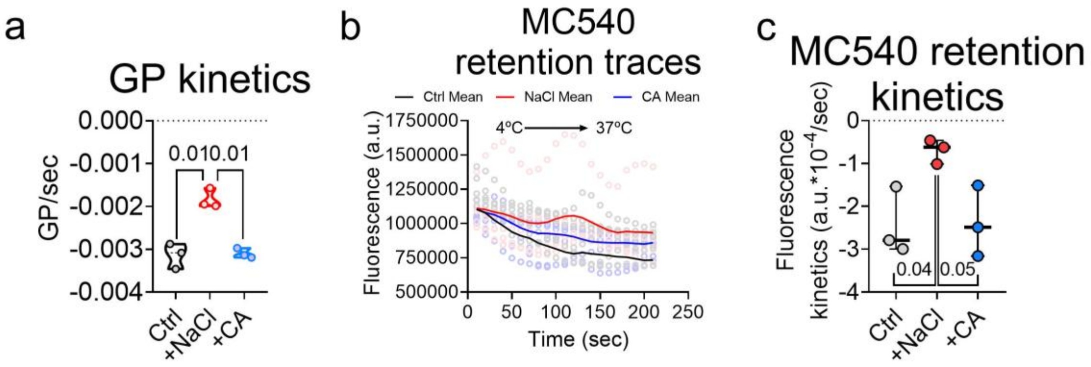


**Extended Data Figure 3. CA does not alter mitochondrial membrane fluidity or phase transition**. **(a)** Fluorescence traces of isolated mitochondrial membranes in which MC540 had been incorporated, treated without (Ctrl Mean) or with 10 mM NaCl (NaCl Mean) or CA (CA Mean). Temperature was used as internal control. Circles represent each time point of each replicate (grey: Ctrl; pink: NaCl; light blue: CAl) (n=3). **(b)** Slope values from data in Fig. 2a. **(c)** Slope values from data in (a). Data are presented as median ± percentiles in (b and c). One-way ANOVA analysis was performed in (b and c).

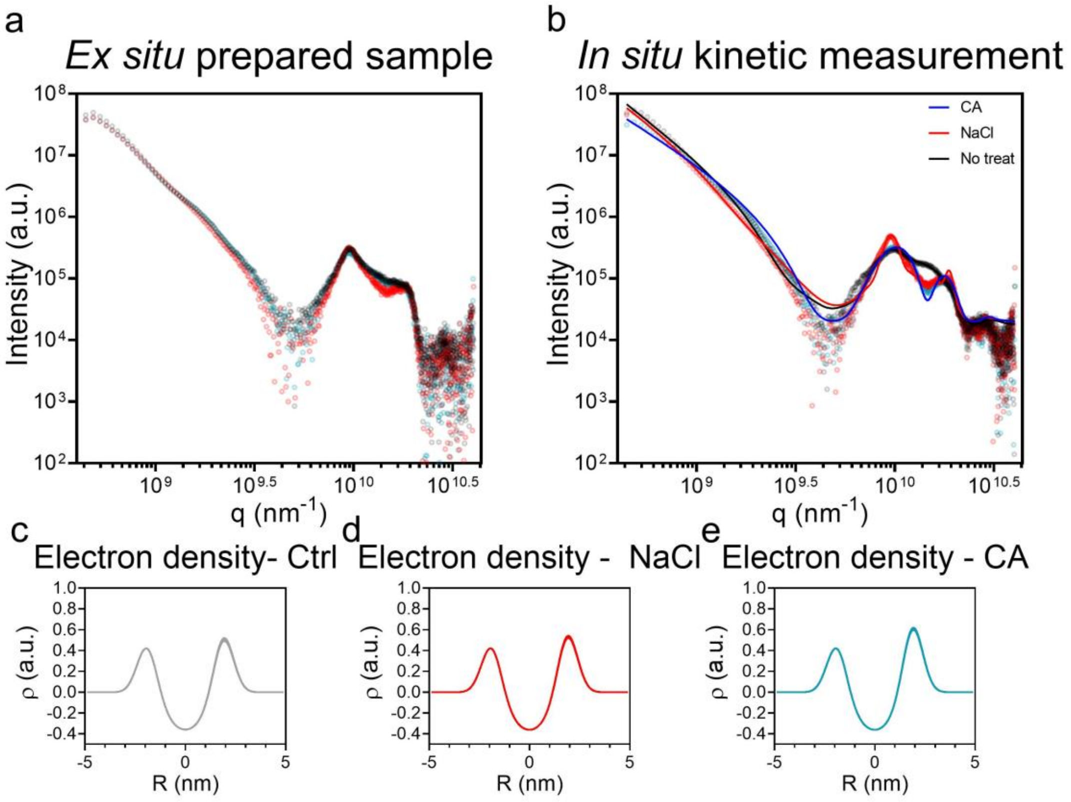


**Extended Data Figure 4. CA interacts with the membrane surface of liposomes**. **(a)** SAXS pattern of *ex situ* prepared samples. **(b)** SAXS pattern of the *in situ kinetic* measurement. *In situ* kinetics is a experimental setup where liposomes and solutions are mixed at the moment of analysis, whereas *ex situ* sample is a sample prepared days before the analysis. **(c-e)** Electron density profiles for ex-situ samples in either untreated liposomes **(c)** or treated with 10 mM NaCl **(d)** or CA **(e)**, measured by Single-Angled X ray Scattering (SAXS)

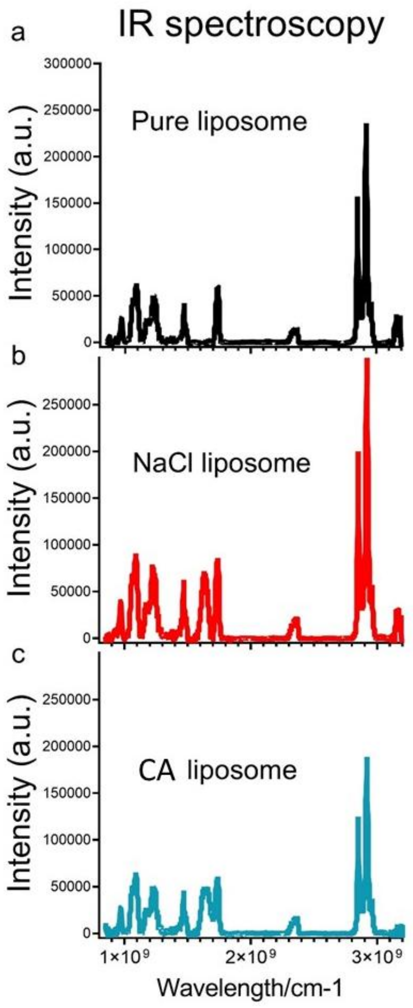


**Extended Data Figure 5. Complete IR spectra of $Na^{+}$ and CA interaction with PC liposomes**. **(a-c)** The entire IR spectra of pure liposomes (a) and liposomes treated with 10 mM NaCl (b) or CA (c) shows that the only peak shifting is the one corresponding to the PC carbonyl group in all cases.

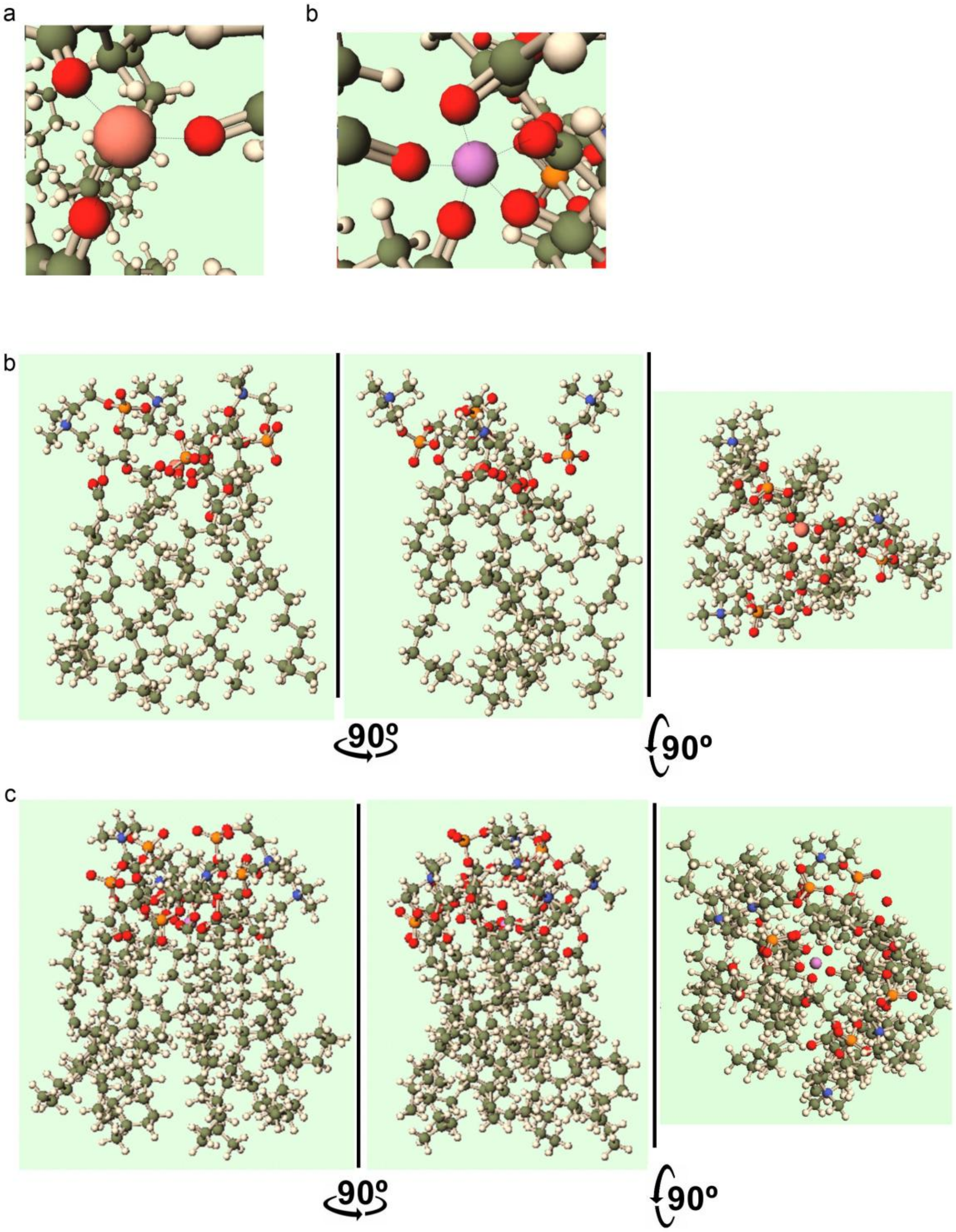


**Extended Data Figure 6. The different geometries in CA:PC and $Na^+$:PC coordinates form adducts with distinct conformations**. **(a)** Molecular representation of the $Na^+$:carbonyl coordination. **(b)** Molecular representation of the CA:carbonyl coordination. **(c)** Molecular representation of the adduct formed after 1$Na^+$:3PC carbonyl coordination. **(d)** Molecular representation of the adduct formed after 1CA:5PC carbonyl coordination. Pink spheres: $Na^+$; Violet spheres: CA; Red spheres: O; Grey spheres: C; White spheres: H; Orange spheres: P; Blue spheres: P.

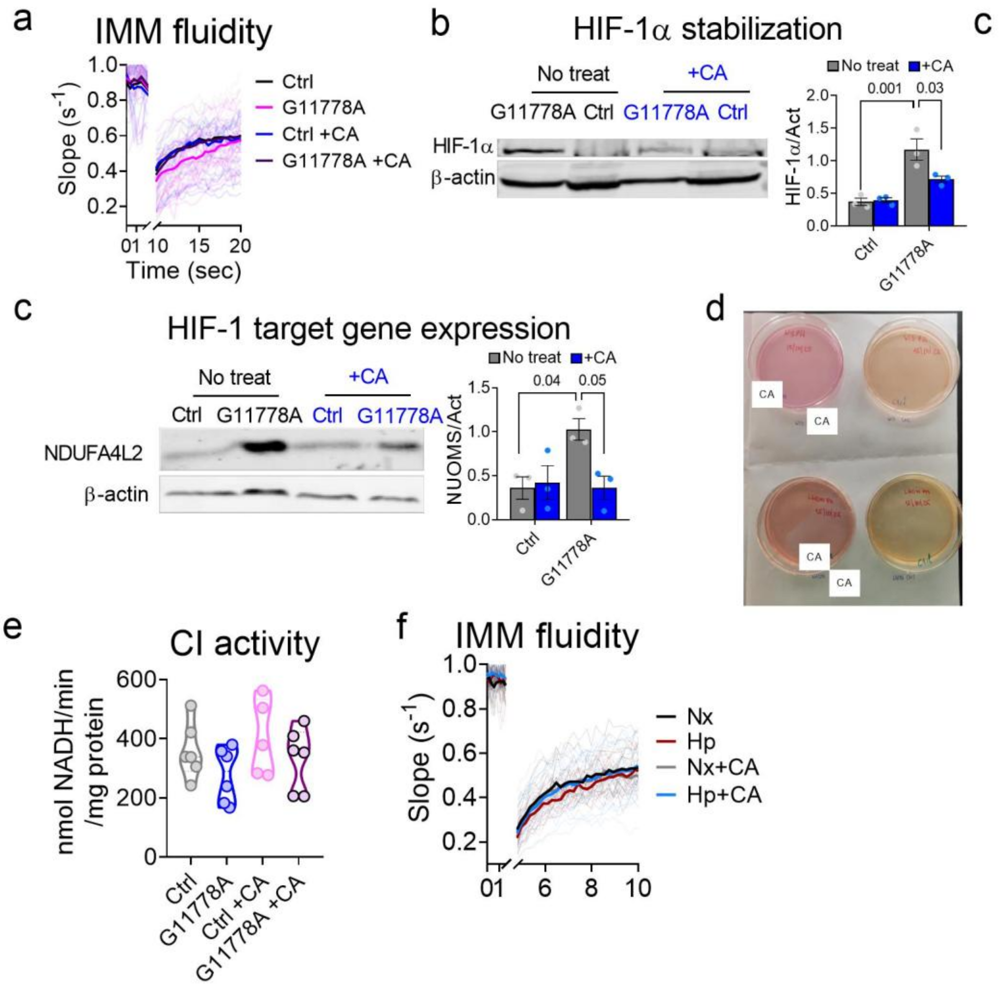


**Extended Data Figure 7. The hypoxic signature in G11778A is reversed by chronic CA treatment**. **(a)** FRAP traces of G11778A and its isogenic control either untreated (Ctrl) or treated with 1 mM CA (n=4 independent experiments; grey: individual Ctrl FRAP traces; light pink: individual G11778A FRAP traces; light blue: individual Ctrl+CA FRAP traces; purple: individual G11778A+CA FRAP traces). **(b)** SDS-PAGE assessing HIF-1α and α-actin levels in G11778A versus Ctrl either untreated or treated with 1 mM CA. Left: representative western blot images; right: quantification of fluorescence. **(c)** SDS-PAGE assessing the HIF-1 target NUOMS and α-actin levels in G11778A versus Ctrl either untreated or treated with 1mM CA. Left: representative western blot images; right: quantification of fluorescence. **(d)** Representative image of culture dishes containing either untreated or 1 mM CA-treated G11778A cells (down) and Ctrl (up) at confluence. **(f)** Rotenone sensitive NADH-driven decylubiquinone reduction of mitochondrial membranes from either untreated or 1mM CA-treated Ctrl or G11778A cells. **(g)** FRAP traces either untreated (Ctrl) or treated with 1 mM CA BAECs that had been subjected to 15 min of noromxia (Nx) or hypoxia (Hp; 1% $O_2$; n=4 independent experiments; grey: individual Ctrl FRAP traces; light pink: individual G11778A FRAP traces; light blue: individual Ctrl+CA FRAP traces; purple: individual G11778A+CA FRAP traces).

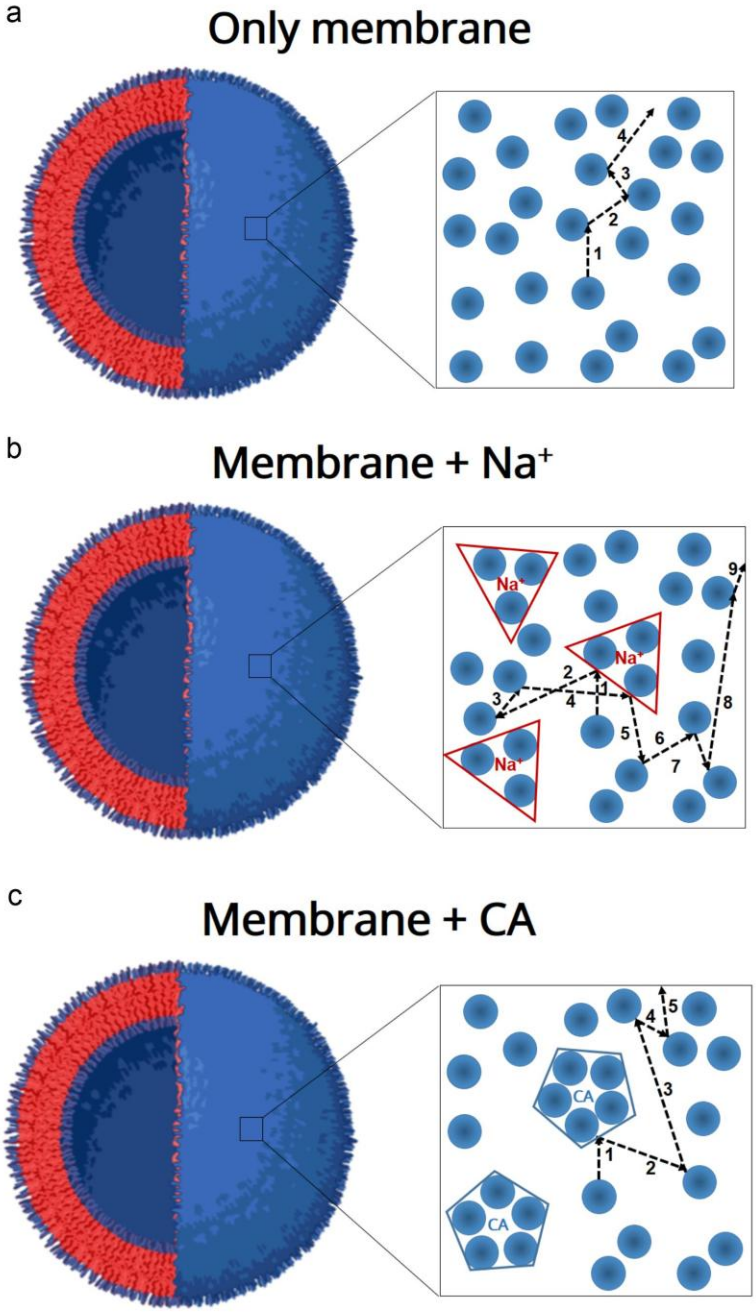


**Extended Data Figure 8. Scheme showing upside down view of lateral phospholipid diffusion in the presence of different single-charged cations**. **(a)** Single phospholipids diffuse freely allowing its rapid advance along the membrane. **(b)** Trigonal $Na^+$:phospholipid adduct impedes free diffusion of single phospholipid molecules, hindering their advance along the membrane. **(c)** Pentameric CA:phospholipid assembly permits free diffusion of single phospholipid molecules due to the less edged nature of the coordinated complex.
